# Evidence supporting a viral origin of the eukaryotic nucleus

**DOI:** 10.1101/679175

**Authors:** Philip JL Bell

## Abstract

The defining feature of the eukaryotic cell is the possession of a nucleus that uncouples transcription from translation. This uncoupling of transcription from translation depends on a complex process employing hundreds of eukaryotic specific genes acting in concert and requires the 7-methylguanylate (m7G) cap to prime eukaryotic mRNA for splicing, nuclear export, and cytoplasmic translation. The origin of this complex system is currently a paradox since it is not found or needed in prokaryotic cells which lack nuclei, yet it was apparently present and fully functional in the Last Eukaryotic Common Ancestor (LECA). According to the Viral Eukaryogenesis (VE) hypothesis the abrupt appearance of the nucleus in the eukaryotic lineage occurred because the nucleus descends from the viral factory of a DNA phage that infected the archaeal ancestor of the eukaryotes. Consequently, the system for uncoupling of transcription from translation in eukaryotes is predicted by the VE hypothesis to be viral in origin. In support of this hypothesis it is shown here that m7G capping apparatus that primes the uncoupling of transcription from translation in eukaryotes is present in viruses of the *Mimiviridae* but absent from bona-fide archaeal relatives of the eukaryotes such as *Lokiarchaeota*. Furthermore, phylogenetic analysis of the m7G capping pathway indicates that eukaryotic nuclei and *Mimiviridae* obtained this pathway from a common ancestral source that predated the origin of LECA. These results support the VE hypothesis and suggest the eukaryotic nucleus and the *Mimiviridae* descend from a common First Eukaryotic Nuclear Ancestor (FENA).

## Introduction

A membrane-bound nucleus defines the eukaryotic domain, and all cellular organisms without nuclei are prokaryotic (Sapp 2005; Stanier and Van Niel 1962). Its presence contributes to a great divide between eukaryotes and prokaryotes defined by features such as linear chromosomes, telomeres, nuclear pores, the spliceosome, mitosis, meiosis, the sexual cycle, and the endoplasmic reticulum. Since the nucleus separates the eukaryotic genome from the ribosomal apparatus, its presence also introduces an uncoupling of transcription from translation unique to the eukaryotic domain.

*Lokiarchaeota* reportedly ‘bridges the gap between prokaryotes and eukaryotes’ and are proposed to be bona-fide archaeal relative of the eukaryotes (Spang et al. 2015). *Lokiarchaeota* and related Asgardians encode Crenactins, the ESCRT-III complex, a family of small Ras-like GTPases and a ubiquitin system, making them a plausible direct descendent of an archaeal ancestor of the eukaryotes (Koonin 2015). This discovery supports the Eocyte tree of life where eukaryotes evolved from a specific archaeon rather than representing a sister group to the archaea (Riviera and Lake 1992). If *Lokiarchaeota* are bona-fide archaeal relatives of the eukaryotes, the last common ancestor of the Asgard archaea and the eukaryotes can be inferred to be an archaeal First Eukaryotic Common Ancestor (FECA) (Eme et al. 2017).

Despite sharing a proposed archaeal ancestor with *Lokiarchaeota*, the last eukaryotic common ancestor (LECA) possessed both a nucleus and a mitochondrion and no eukaryotes are descended from any earlier intermediates without both these complex organelles (Neumann et al. 2010). Whilst the abrupt appearance of the mitochondrion in LECA is persuasively explained by its endosymbiotic descent from an alpha-proteobacterium (e.g. Lang et al. 1999) the similarly abrupt appearance of a nucleus has been much more difficult to explain (Martin 1999; Martin 2005).

The presence of the eukaryotic nucleus results in an uncoupling transcription and translation, and this uncoupling requires mRNA to be synthesised inside the nucleus, capped, processed, and exported into the cytoplasm for translation (Kyrieleis et al. 2014). This contrasts to prokaryotic translational systems that rely on direct recognition of uncapped mRNA by the ribosomal apparatus (Benelli and Londei 2011). Evidence of archaeal methanogens 3.8 billion years ago (Battistuzzi et al. 2004) shows that prokaryotes evolved well before the eukaryotes originated 1.8 billion years ago (Parfrey et al. 2011). Accordingly the prokaryotic system of direct recognition of mRNA by the ribosomal apparatus existed for nearly two billion years before the nucleus and its cap based system abruptly appeared in LECA.

The change from a prokaryotic translational system found in FECA (**Figure 1**) to the uncoupled eukaryotic system found in LECA (**Figure 2**) involved the evolution of a complex molecular system involving hundreds of interacting genes. The m7G cap is critical to this process since it primes the mRNA for processing, export and translation (**Figure 2**). The genes required to add the m7G cap include: an RNA polymerase (RNAP-II) dedicated to capped mRNA synthesis (Sentenac 1985), an RNA triphophatase (TPase), guanylyltransferase (GTase) and methyltransferase (MTase) required for capping mRNA (Kyrieleis et al. 2014). A cap binding protein (eIF4E) is also essential since it is required for initiating translation of the capped mRNA in the cytoplasm (Marcotrigiano et al. 1997). Paradoxically, the high level of complexity and the integrated nature of the cap based system of uncoupling transcription from translation suggest a long evolutionary history, yet no transitional cellular forms linking the prokaryotic (**Figure 1**) and eukaryotic systems (**Figure 2**) have been described. Consequently if only prokaryotes are considered as source for the eukaryotic m7G cap based system, an abrupt and currently insurmountable phylogenetic impasse is encountered.

**Figure 1:**
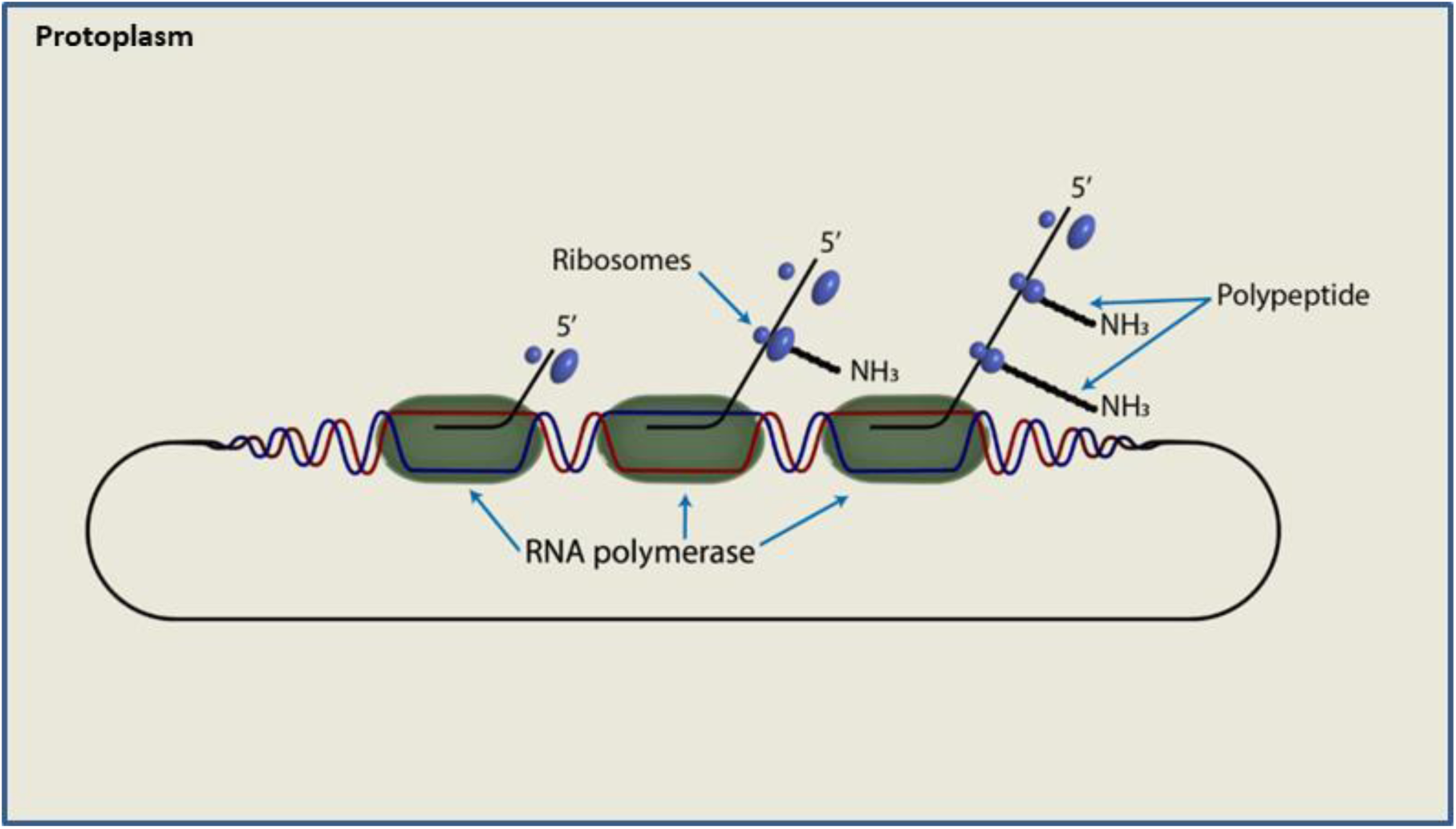
The coupled prokaryotic system of transcription and translation. Both archaea and bacteria utilize one type of multi-component RNA polymerase (RNAP) to transcribe all RNA (Werner 2007). Transcription and translation in prokaryotes are coupled since transcription and translation occur directly in the protoplasm, and thus translation initiation can occur before the mRNA transcript is fully synthesised. Translation in prokaryotes relies on direct recognition of mRNA by the ribosomal apparatus via sequences such as the Shine-Dalgarno sequences or short UTR’s (Benelli and Londei 2011). In the case of Shine-Dalgarno sites, the 30S ribosomal subunit binds to the mRNA in such a way that AUG codon lies on the peptidyl (P) site and the second codon lies on aminoacyl (A) site. The initiator tRNA binds to the P site, the large ribosomal subunit docks with the small subunit, the initiation factors are released and the ribosome is ready to start translation. Since prokaryotes originated 3.8 billion years ago (Battistuzzi et al. 2004) the coupled prokaryotic process predates the uncoupled eukaryotic system by close to 2 billion years and thus is the most ancient cellular system of transcription and translation.

**Figure 2:**
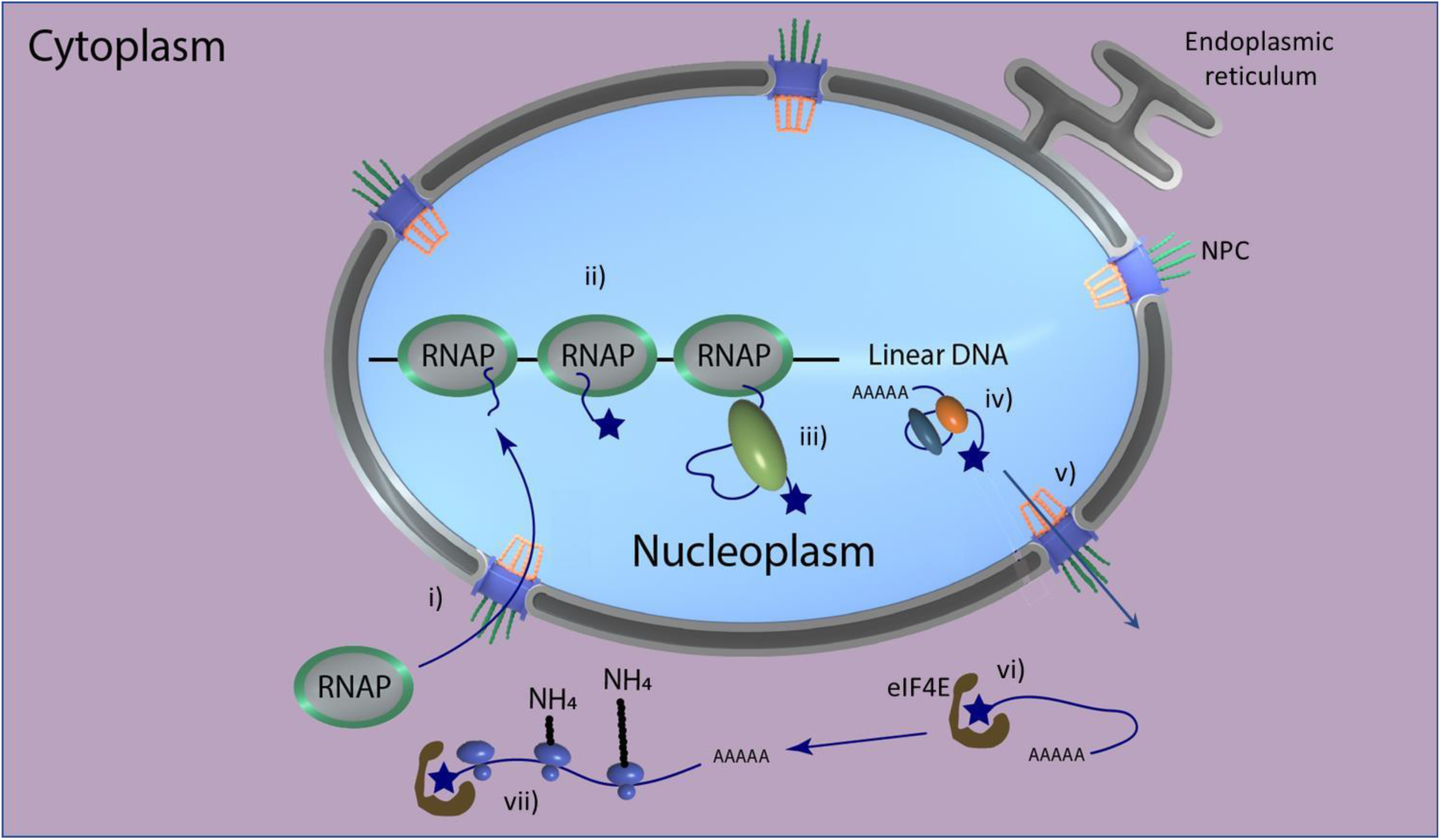
The eukaryotic system to uncouple transcription from translation is complex and employs hundreds of genes that act in concert. The dominant eukaryotic cap dependent system of transcription and translation that was apparently present and fully functional in LECA (Neumann et al. 2010) is described below. Subunits of RNAP-II are translated in the cytoplasm imported into the nucleus through the Nuclear Pore Complex (NPC). RNAP-II initiates transcription of mRNA by binding to the promoter regions of protein coding genes. **ii)** After the synthesis of the first 20 to 25 bp of mRNA, the polymerase pauses until the mRNA is capped (Ramanathan et al. 2016, Okamura et al. 2015). The eukaryotic m7G cap (symbolised by ✶) consists of 7-methylguanosine linked via a reversed 5’-5’ triphosphate linkage to the transcript and is the first modification made to RNAP-II transcribed RNA. Three enzymatic functions are required to generate the cap. Firstly, a RNA 5’-phosphatase (TPase) hydrolyses the 5’-triphosphatase end of the nascent mRNA to generate a 5’-diphosphate. The 5’-diphosphate is then capped with guanosine mono-phosphate by a RNA guanylyltransferase (GTase) to generate a 5’ GpppRNA cap on the transcript. Finally, the guanosine GpppRNA cap is methylated by RNA (guanine-N7)-methyltransferase (MTase) (Kyrieleis et al. 2014). **iii)** The nuclear cap binding complex (CBC) binds to the m7G cap which then forms a complex with snRNP’s to initiate splicing and polyadenylation (Ramanathan et al. 2016). Splicing of mRNA transcripts is unique to the eukaryotes and requires interaction of hundreds of proteins and the conserved snRNAs. **iv)** The m7G cap primes the mRNA for transport through the nuclear pores into the cytoplasm (Katahira 2015) by binding trans-acting factors to form a mature messenger ribonucleoprotein (mRNPs). Recruitment of the multisubunit TRanscription-EXport (TREX-complex requires the 5’ capping of pre-mRNA because CBP80 interacts with the QAlyRef and THO sub-complexes of TREX-1 (Okamura et al. 2015). **v)** The nuclear pore complex (NPC) is integral to the uncoupling of transcription from translation because the NPC acts as a gate keeper, controlling which macromolecules enter and exit the nucleus. NPC’s are unique to the eukaryotes, and a single NPC comprises ∼500 individual protein molecules collectively known as nucleoporins (Nups) (Kabachinski and Schwartz 2015). The NPC includes a nuclear ring, a central transport channel and eight cytoplasmic fibrils which allow molecules smaller that 40-60 kDa to freely diffuse (Kabachinski and Schwartz 2015). Large molecules such as mRNA must associate with specific export receptors such as Nxf1, Crm1 or other karyopherins to be actively transported through the NPC. **vi)** To initiate translation, the 43S ribosomal preinitiation complex is recruited to the 5’ end of the mRNA, a process that is co-ordinated by eIF4E through its interactions with eIF4G and the 40S ribosomal subunit associated eIF3 (Hernandez et al. 2012). Several eukaryotic specific initiation factors eIF4E, eIF4G, eIF4B, eIF4H and eIF3 are involved with 5’-cap-binding and scanning processes that are essential to the initiation and translation of capped eukaryotic mRNA (Jagus et al. 2012). **vii)** Once the ribosome has been recruited to the capped mRNA transcript, a scanning process occurs and translation is generally initiated at the first ATG encountered.

The Viral Eukaryogenesis (VE) hypothesis proposes the nucleus derives from an ancient DNA phage/virus and predicts the m7G cap based system that primes the uncoupling of transcription from translation in eukaryotes originated amongst the prokaryotic viruses (Bell 2001). Although fossil evidence for viruses is unlikely to be found, viruses almost certainly existed before the origin of LECA. For example, a prokaryotic genome that is free of genetic parasites is expected to show signs of genome degeneration due to the need for a mechanism to overcome the degradation of prokaryotic genomes caused by processes such as Muller’s ratchet (Iranzo et al. 2016). There are also strong biological arguments that the emergence of genetic parasites is inevitable due to the instability of parasite-free states (Koonin et al. 2017). Further experimental support for a pre-LECA origin of viruses comes from phylogenomic analysis which shows that modern eukaryotic viruses evolved from pre-existing prokaryotic phage (Koonin et al. 2015). It can thus be anticipated that viruses would have emerged in concert with the first prokaryotes and existed for much of the 2 billion years between the appearance of the first methanogens and the appearance of LECA.

The VE hypothesis has been supported by the discovery that the *Pseudomonas* jumbophage *201 Φ2-1* constructs a nucleus-like viral factory that uncouples transcription from translation (Chaikeeratisak et al. 2017b). The viral factory established by *201 Φ2-1* confines phage DNA within the factory whilst excluding ribosomes (Chaikeeratisak et al. 2017b). Thus once the factory is established, transcription occurs within the factory and the mRNA must be exported into the cytoplasm for translation. Functionally, infection results in the bacterial protoplasm being divided into a viroplasm where viral information processing occurs, and a cytoplasm where translation and metabolic enzymes are localised. Since viral encoded enzymes such as RNA polymerases and DNA polymerases must function inside the viral factory whilst components of the phage virions are assembled in the cytoplasm, it can be inferred that the boundary of these viral factories must be able to selectively sort which proteins, RNA transcripts and other factors can move across the boundary.

Deepening similarities between the eukaryotic nucleus and the viral factories of phage *201 Φ2-1, 201 Φ2-1* possesses homologues of eukaryotic tubulin (*PhuZ*), and this tubulin polymerises via dynamic instability, positioning the factory in the centre of the infected cell (Chaikeeratisak et al. 2017). The *PhuZ* spindle is the only known example of a cytoskeletal structure that shares three key properties with the eukaryotic spindle: dynamic instability, a bipolar array of filaments, and central positioning of DNA (Chaikeeratisak et al. 2017). This is a significant parallel since eukaryotic nuclei are positioned in the cell by microtubule-dependent motors during development and differentiation (Star 2009).

It was similarities between the eukaryotic nucleus and the Pox viruses that led to the original VE proposal that the nucleus was derived from a virus that infected the archaeal ancestor of the eukaryotes (Bell 2001). In particular the observations supporting the model were that the Pox viruses could produce capped mRNA, possessed linear chromosomes, could separate transcription from translation, and had an ability to replicate entirely within the host cytoplasm (Bell 2001). Subsequently Pox viruses were found to be members of an ancient monophyletic group, the NCLDV viruses (Iyer et al. 2001). The discovery of the giant Mimivirus in 2004 and its allocation to the NCLDV group (Raoult et al. 2004) demonstrated pox-viral relatives were of unprecedented size and possessed a complexity comparable to prokaryotic cells (Raoult et al. 2004). Many other giant NCLDV viruses have been discovered including even more complex relatives such as the *Kloseneuvirus* (Schultz et al. 2017) and *Tupanvirus* (Abrahão et al. 2018).

A prokaryotic viral ancestry for both the Poxviruses and the other NCLDV viruses has been supported by phylogenomic studies (Koonin and Yutin 2010) and is compatible with the NCLDV common ancestor existing at or before the origin of LECA (Boyer et al. 2010; Nasir et al. 2012; Yutin et al. 2009). Furthermore, comparison between inferred genome of the NCLDV common ancestor (Yutin and Koonin 2012) and the modern *PhiKZ* like viruses (including *201 Φ2-1*) reveals that both classes of giant virus possess large genomes, encode homologues of DNA polymerases (Kazlauskas and Venclovas 2011), multi-subunit RNA polymerase (Ceyssens et al. 2014), DNA ligases (Wojtus et al. 2017), RNA ligases (Wojtus et al. 2017) and replicate with a high degree of autonomy from their hosts (Yuan and Gao 2017). Furthermore distantly related NCLDV viruses from the *Poxviridae, Asfaviridae, Pithoviridae Marseilleviridae and Mimiviridae* families replicate partially or exclusively within large cytoplasmic Viral Factories (Fridmann-Sirkis 2016). Since the genes for adding an m7G cap to mRNA were present in the common ancestor of all the NCLDV viruses (Iyer et al. 2001; Yutin and Koonin 2012) it can be inferred from these observations that the common ancestor of the NCLDV viruses was a virus that could produce capped mRNA and like phage *201 Φ2-1* could establish a viral factory in its host’s cytoplasm.

In addition to inheriting the ability to add an m7G cap to mRNA from the NCLDV common ancestor, two separate groups of NCLDV viruses, the *Pandoraviridae* and the *Mimiviridae*, also possess homologues of the eukaryotic cap binding protein eIF4E (Schultz et al. 2017). Unlike many NCLDV viruses, the Pandoraviruses possess introns in their genes strongly suggesting that at least part of the *Pandoravirus* genome is transcribed in the nucleus (Phillipe et al. 2013). By contrast members of the *Mimiviridae* have been shown to replicate entirely in the host cytoplasm and establish a nucleus-like uncoupling of transcription from translation (Fridmann-Sirkis et al. 2016). Furthermore, the cap binding protein encoded by eIF4E is located in the cytoplasm outside the viral factory during infection (Fridmann-Sirkis et al. 2016). Thus as shown in **Figure 3**, in addition to viral factories of Mimiviruses and *201 Φ2-1* sharing fundamental features with each other such as the ability to uncouple transcription from translation and selectively control which macromolecules enter and exit the viral factory, the Mimivirus viral factories also share further specific fundamental features with the eukaryotic nucleus. Amongst the shared features with the eukaryotic nucleus, the Mimivirus has a linear genome, establishes a nucleus-like organelle in its host’s cytoplasm, possesses its own RNA polymerase dedicated to transcribing capped mRNA, exports capped mRNA into the cytoplasm for translation, and possesses its own version of the cap binding protein (eIF4E) which is located in the host cytoplasm during infection and is presumably involved in controlling the initiation of translation of capped Mimiviral transcripts. These discoveries have led to independent suggestions that the nucleus is a derived from a viral factory (e.g. Forterre and Raoult 2017).

**Figure 3:**
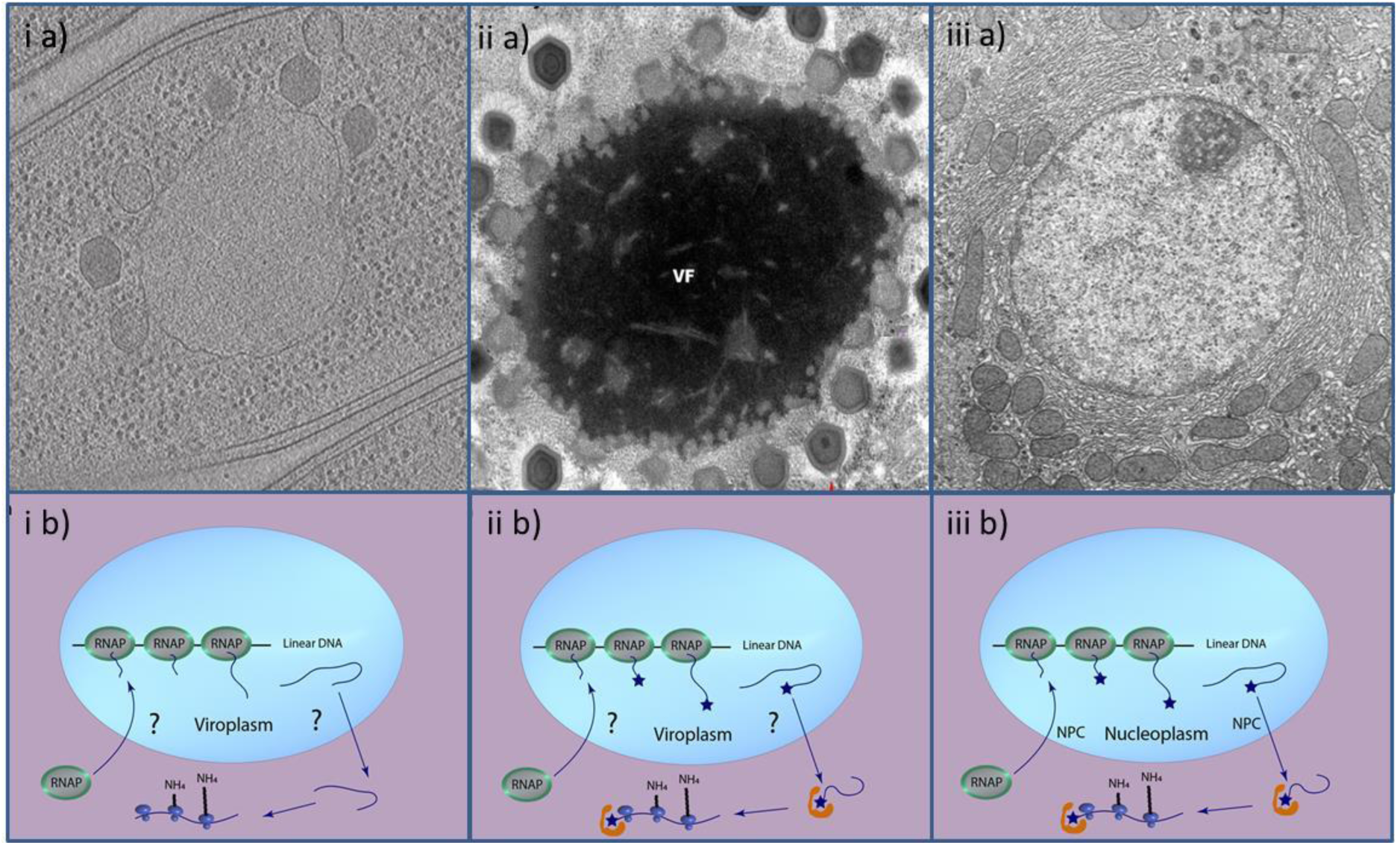
*201 Φ2-1* viral factories, Mimivirus viral factories, and the eukaryotic nucleus share the ability to uncouple transcription from translation. **i a)** Image of Phage *201 Φ2-1* viral factory (Chaikeeratisak et al. 2017b). **ii a)** Image of Mimivirus viral factory (Zauberman et al. 2008). **iii a)** The eukaryotic nucleus. **i b)** Phage *201 Φ2-1* establishes a viral factory in the cytoplasm of the bacterial host confining DNA replication and transcription to the viral factory. Translation is confined to the cytoplasm since host bacterial ribosomes are excluded from the viral factory (Chaikeeratisak et al. 2017b). Since *PhiKZ* relatives of *201 Φ2-1* can complete infection in the absence of bacterial RNA polymerase (RNAP) activity (Ceyssens et al. 2014) it can be inferred that the multi-subunit RNAP genes encoded by the phage are transcribed in the viral factory, transcripts exported into the cytoplasm for translation and the proteins re-imported into the viral factory to transcribe the phage DNA. **ii b)** The Mimivirus also establishes a viral factory in the cytoplasm of its eukaryotic host (Mutsafi et al. 2010) confining DNA replication and transcription to the viral factory. Translation is confined to the cytoplasm since host ribosomes are excluded from the viral factory (Fridmann-Sirkis et al. 2016). Mimiviruses encode a multi-subunit RNA polymerase that transcribes Mimiviral DNA and functions within the viral factory (Fridmann-Sirkis et al. 2016). It can therefore be inferred that the Mimivirus viral factory controls which macromolecules are transported in and out of the viroplasm. Like the eukaryotic nucleus *Mimiviridae* encode their own mRNA capping apparatus and a version of the eIF4E gene. In cells infected by the *Mimiviridae* EIF4E remains located in the host cytoplasm (Fridmann-Sirkis et al. 2016). **iii b)** The eukaryotic nucleus, like viral factories of both phage *201 Φ2-1* and the Mimivirus, is a specialised compartment located in the cytoplasm that confines DNA replication and transcription within its boundaries. Translation is confined to the cytoplasm since functional ribosomes are excluded from the nucleus. The mRNA encoding RNAP-II subunits are transcribed within the nucleus, exported into the cytoplasm for translation, and re-imported into the nucleus to transcribe nuclear DNA. Unlike viral factories, the mechanisms by which the nucleus sorts the macromolecules that can enter and exit the nucleus is well understood, and known to be controlled by the NPCs. Like the Mimivirus viral factory, eukaryotic nuclei encode their own capping apparatus and encode the eIF4E gene which binds to the m7G cap and both are part of a complex system to uncouple transcription from translation.

Amongst the phylogenetically diverse NCLDV viruses, the *Mimiviridae* appear to be the only group that establishes a solely cytoplasmic viral factory and possesses the eIF4E gene. Thus by analogy with the proposal that the presence of Crenactins, the ESCRT-III complex, a family of small Ras-like GTPases and a ubiquitin system make *Lokiarchaeota* a plausible direct descendent of an archaeal ancestor of the eukaryotes (Koonin 2015) it is proposed here that ability to construct a viral factory that uncouples transcription from translation, the possession of the m7G capping apparatus and the presence of the eIF4E binding protein make the Mimiviridae a plausible direct descendant of a viral ancestor of the eukaryotic nucleus. It is proposed here that this common ancestor of the *Mimiviridae* and the eukaryotic nucleus was the First Eukaryotic Nuclear Ancestor (FENA). To test this hypothesis, phylogenetic analysis was performed on the largest subunit of RNAP-II which is required for synthesis of mRNA destined for capping; the capping apparatus which are required to add the m7G cap to eukaryotic mRNA, and the eIF4E gene which is required to initiate translation of capped mRNA in the cytoplasm (see **Figure 2**).

## Results

### The closest known archaeal relative of the eukaryotes shows no evidence of the eukaryotic genes required to prime the uncoupling of transcription from translation in eukaryotes

To confirm the extent to which the closest archaeal relatives of the eukaryotes lack homologues of the eukaryotic cap based system for uncoupling of transcription from translation, the genome of *Heimdallarchaea LC-3* (formerly Loki3, (Spang et al. 2015; Spang et al. 2018)) was Blast searched to identify homologues of four of the *Saccharomyces cerevisiae* genes required to uncouple transcription from translation.

*RPO21* of *S. cerevisiae* was used to identify homologues of RNAP-II in *Heimdallarchaea LC-3. RPO21* was chosen because it is the largest subunit of RNAP-II in *S. cerevisiae*, and it encodes the C Terminal Domain (CTD) containing the heptapeptide repeat (YSPTSPS) that recruits the capping enzymes to the nascent mRNA transcript and is intricately involved in further processing of capped mRNA (McCracken et al. 1997). The *Homo sapiens* genome was also searched for homologues to illustrate the level of homology of these genes in distantly related descendants of LECA. Despite the large evolutionary distance between the yeast and humans, three different homologues of *RPO21* with very significant E values were identified in *H. sapiens* (**Table 1**). These three correspond to RNA-I, RNA-II and RNA-III which are are present in all eukaryotes (Sentenac 1985). By contrast in *Heimdallarchaeota LC-3*, only a single RNA polymerase subunit A’ was identified as a homologue. This is consistent with *Heimdallarchaoeta LC-3* possessing a prokaryotic transcription system where all RNA is transcribed by the same RNA polymerase (Werner 2007).

**Table 1.** Homologues of *S. cerevisiae* RNAP-II, GTase, MTase and eIF4E in *Homo sapiens* and *Heimdallarchaota LC-3* identified using Blast.

| Saccharomyces gene name | Annotation in *Homo sapiens* | *Homo sapiens* |  | *Heimdallarchaeota LC-3* |  | Annotation in *Heimdallarchaeota LC-3* |
| --- | --- | --- | --- | --- | --- | --- |
|  |  | Accession | E value | Accession | E value |  |
| RPO21 (NP_010141) | RNA polymerase I large subunit | NP_056240 | 1.00E-97 |  |  |  |
|  | RNA polymerase II large subunit | NP_000928 | 0 | OLS19521.1 | 0 | RNA polymerase subunit A' |
|  | RNA polymerase III large subunit | NP_008986 | 0 |  |  |  |
| CEG1 (NP_011385) | Guanylyltransferase | AAH19954 | 2.00E-17 | OLS26805.1 | 2.7 | DNA ligase |
| ABD1 (NP_009795) | Methyltransferase | NP_003790 | 2.00E-38 | OLS26405.1 | 4.00E-05 | Trans-aconitate 2-methyltransferase |
| CDC33 (NP_014502) | eIF4E | NP_001959 | 3.00E-36 | OLS27732.1 | 0.37 | 5-exo-hydroxycamphor dehydrogenase |

**Table 2.** Summary of Accession numbers used in this study

| Group | species | Gtase | Mtase | eIF4E | RNAP-II | RNAP-III |
| --- | --- | --- | --- | --- | --- | --- |
| Eukarya Fungi | *Saccharomyces cerevisiae* | NP_011385 | NP_009795 | NP_014502 | NP_010141 | NP_014759 |
| Eukarya Fungi | *Kluyveromyces marxianus* | XP_022676394 | XP_022674569 | XP_022678436 | XP_022677581 | XP_022678447 |
| Eukarya Fungi | *Aspergillus niger* | XP_001400555 | XP_001394253 | XP_001395221 | XP_001389676 | XP_001393726 |
| Eukarya Holozoa | *Homo sapiens* | AAH19954 | BAA82447 | NP_001959 | NP_000928 | NP_008986 |
| Eukarya Holozoa | *Mus musculus* | NP_036014 | NP_080716 | NP_031943 | AAB58418 | NP_001074716 |
| Eukarya Holozoa | *Danio rerio* | NP_998032 | NP_001038465 | NP_001007778 | XP_005156282 | NP_001263425 |
| Eukarya Holozoa | *Caenorhabditis elegans* | NP_001020979 | NP_492674 | NP_503124 | NP_500523 | NP_501127 |
| Eukarya viridiplantae | *Arabidopsis lyrata* | XP_002873017 | XP_002894293 | XP_020875354 | XP_020873010 | XP_020884300 |
| Eukarya viridiplantae | *Brassica napus* | XP_013647283 | XP_013640697 | AGA20262 | XP_013656472 | XP_013681133 |
| Eukarya viridiplantae | *Ostreococcus tauri* | XP_003075327 | XP_003081423 | XP_022840751 | XP_022839775 | XP_022840814 |
| Eukarya Amoebozoa | *Dictyostelium discoideum* | XP_636333 | XP_642389 | XP_647593 | XP_641735 | XP_642724 |
| Eukarya Amoebozoa | *Dictyostelium purpureum* | XP_003293052 | XP_003293647 | XP_003293106 | XP_003285719 | XP_003284018 |
| Eukarya Amoebozoa | *Acytostelium subglobosum LB1* | XP_012756463 | XP_012752660 | XP_012756585 | XP_012756853 | XP_012752065 |
| Eukarya Alveolata | *Plasmodium falciparum* | KNC37820 | ETW19449 | XP_001351220 | XP_001351252 | XP_001350009 |
| Eukarya Alveolata | *Plasmodium vivax* | KMZ83875 | SGX75114 | XP_001614562 | XP_001614530 | XP_001614080 |
| Eukarya Alveolata | *Theileria equi strain WA* | XP_004828897 | XP_004828862 | XP_004829399 | XP_004831990 | XP_004830926 |
| Eukarya Alveolata | *Perkinsus marinus ATCC 50983* | XP_002774114 | XP_002774250 | XP_002774365 | XP_002767562 | XP_002778409 |
| Eukarya Alveolata | *Cryptosporidium muris RN66* | XP_002140608 | XP_002139632 | XP_002140059 | XP_002141559 | XP_002142344 |
| Eukarya Excavata | *Trypanosoma cruzi cruzi* | PBJ71163 | PBJ71163 | PBJ73557 | PBJ81421 | PBJ72541 |
| Eukarya Excavata | *Trypanosoma rangeli* | RNF00410 | RNF00410 | RNF02202 | RNF07318 | RNF04215 |
| Eukarya Excavata | *Leishmania mexicana* | XP_003875466 | XP_003875466 | XP_003876737 | XP_003877779 | XP_003878621 |
| Eukarya Excavata | *Leptomonas seymouri* | KPI83387 | KPI83387 | KPI89876 | KPI84927 | KPI86235 |
| Eukarya Excavata | *Bodo saltans* | CUG90421 | CUG90421 | CUF95139 | CUI14899 | CUI14455 |
| Mimiviridae Klosneuvirinae | *Catovirus CTV1* | ARF09224 | ARF09224 | ARF09024 | ARF09013-20 |  |
| Mimiviridae Klosneuvirinae | *Klosneuvirus KNV1* | ARF11732 | ARF11732 | ARF11337 | ARF11340-43 |  |
| Mimiviridae Klosneuvirinae | *Indivirus ILV1* | ARF09638 | ARF09638 | ARF09452 | ARF09455 |  |
| Mimiviridae Klosneuvirinae | *Bodo saltans virus* | ATZ80933 | ATZ80933 | ATZ80516 | ATZ80519 |  |
| Mimiviridae Mimivirinae | *Acanthamoeba polyphaga mimivirus* | AEJ34618 | AEJ34618 | AKI79272 | YP_003987013 |  |
| Mimiviridae Mimivirinae | *Moumouvirus australiensis* | AVL94825 | AVL94825 | AVL94704 | AVL94698 AVL94696 |  |
| Mimiviridae Mimivirinae | *Powai lake megavirus* | ANB50623 | ANB50623 | ANB50499 | ANB50494 ANB50492 |  |
| Mimiviridae Mimivirinae | *Tupanvirus deep ocean* | AUL79325 | AUL79325 | AUL79602 | AUL79608 |  |
| Mimiviridae Mimivirinae | *Tupanvirus soda lake* | AUL78031 | AUL78031 | AUL78296 | AUL78302 |  |
| Mimiviridae Mimivirinae | *Acanthamoeba polyphaga moumouvirus* | YP_007354410 | YP_007354410 | YP_007354285 | YP_007354277 |  |
| Mimiviridae Mesomimivirinae | *Chrysochromulina ericina virus* | YP_009173557 | YP_009173557 | YP_009173322 | YP_009173653 |  |
| Mimiviridae Mesomimivirinae | *Phaeocystis globosa virus* | YP_008052553 | YP_008052553 | YP_008052407 | YP_008052581 |  |
| Mimiviridae Mesomimivirinae | *Tetraselmis virus 1* | AUF82182 | AUF82182 | AUF82209 | AUF82600 |  |
| Mimiviridae CroV | *Cafeteria roenbergensis virus BV-PW1* | YP_003969844 | YP_003969844 | YP_003969852 | YP_003970001 |  |
| Archaea Crenarchaeota | *Saccharolobus solfataricus* |  |  |  |  | WP_009990476 |
| Archaea Crenarchaeota | *Sulfolobus acidocaldarius* |  |  |  |  | WP_011277574 |
| Archaea Asgardarchaea | *Candidatus Odinarchaeota archaeon LCB_4* |  |  |  |  | OLS17382 |
| Archaea Euryarchaeota | *Pyrococcus furiosus* |  |  |  |  | WP_014835440 |

The capping apparatus in eukaryotes requires a TPase, a GTase and an MTase. Since the TPase gene in eukaryotes apparently arose from two phylogenetically different origins (Ramanathan et al. 2016; Kyrieleis et al. 2014), only the GTase and MTase genes required for constructing the m7G cap were used to search the *Heimdallarchaea LC-3* genome. Using the *CEG1* (GTase) of *S. cerevisiae* to search for homologues in *H. sapiens* identifies the human GTase with a very significant E value. By contrast, although some putative homologues with non-significant E values were identified with Blast in *Heimdallarchaeota LC-3*, only one of these showed homology with the known domain structure of the GTase. This gene was an ATP Ligase (**Table 1**), a group known to share homology with the GTases (Shuman and Schwer 1995). Using the *ABD1* (MTase) gene of *S. cerevisiae* identifies homologs with significant E values in both humans and *Heimdallarchaeota LC-3*. However it is known that the methyltransferase domain of the capping enzyme shares homology with a wide family of methyltransferases, and according to the annotated genome of the *Heimdallarchaeota LC3*, the gene detected in this search shares affinity with the Trans-aconitate 2-methytransferases rather than the capping MTase.

Using the *S. cerevisiae* eIF4E gene (*CDC33*) to search for homologues in *H. sapiens* identifies the human eIF4E with a very significant E value (**Table 1**). By contrast, no homologues eIF4E with significant E values were found in the *Heimdallarchaeaota LC-3* genome. Furthermore, none of the genes with even low degrees of homology detected in *Heimdallarchaeota LC-3* possessed the conserved sites that are known to be involved when eIF4E binds the m7G cap (Marcotrigiano et al. 1997).

These results are consistent with the Asgard archaea being authentically archaeal in design and thus lacking a nucleus, the defining feature of the eukaryotic domain (Stanier and Van Niel 1962). The absence of any sign of a nucleus in Asgard archaea and the sudden appearance of the nucleus in LECA is strikingly similar to the sudden appearance of the mitochondrion in the eukaryotic lineage. Due to fundamental similarities between the mitochondria and alpha-proteobacteria, the abrupt appearance of the mitochondrion in LECA is widely accepted to be the result of endosymbiosis between a bacterium and the ancestor of the eukaryotes (e.g. Lang et al. 1999). The similar abrupt appearance of a highly complex nucleus in LECA in consistent with an endosymbiotic origin, but the nucleus is clearly not of prokaryotic cellular origin since it lacks an obvious homologue or precursor among prokaryotes and is primarily an information processing organelle (Martin 1999; Martin 2005).

### Mimiviral and eukaryotic RNAP-II, Gtase, MTase and eIF4E form two discrete monophyletic groups

Unlike any known prokaryotes, members of the *Mimiviridae* construct nucleus-like viral factories in the cytoplasm of their hosts that separate transcription from translation (**Figure 3**). They also possess a functional mRNA capping pathway that is homologous to the pathway utilised by the eukaryotes to prime the uncoupling of transcription from translation. This pathway includes an RNAP dedicated to mRNA synthesis, a TPase, GTase and MTase required to add the m7G cap to the mRNA, and eIF4E, the cytoplasmic cap binding protein required to initiate translation of the capped transcript in the cytoplasm.

In this phylogenetic analysis members of the eukaryotic domains were carefully selected to cover the major eukaryotic supergroups and thus span the diversity of the eukaryotic domain (see Materials and Methods). Where possible eukaryotic clades were chosen that contained at least one member that has been studied in depth at a molecular level and where experimental knowledge of the processes of transcription and translation exists. In addition, all phylogenetic analysis uses the same organisms, and only species with complete genomes where all genes (RNAP-II, GTase, MTase and eIF4E) could be unambiguously identified were used in tree construction.

As shown **Figure 4a** the unrooted phylogenetic tree of the RNAP largest subunit resolves into two discrete monophyletic clades: the eukaryotes which descend from LECA, and the *Mimiviridae* that descend from the common ancestor of the *Mimiviridae*. Despite the more limited phylogenetic information contained in the GTase, MTase and eIF4E alignments, similar patterns are observed to the RNAP tree, and the monophyly of the eukaryotes and the *Mimiviridae* is maintained in each case. Concatenating the four genes (Fig. 4e) generates a phylogenetic tree with bootstrap values higher than any of the individual trees suggesting that the four genes have a common phylogenetic signal. Within the eukaryotic domain, clades corresponding to *Holozoa, Ameobozoa, Fungi, Viridiplantae, Alveolata* and *Excavata* were well resolved with high support. These results are consistent with studies that show LECA possessed a functional eukaryotic nucleus (Neumann et al. 2010) and that all four eukaryotic genes identified as critical in uncoupling transcription from translation primed by the m7G cap descend from a common ancestral set of genes that were present in LECA. The *Mimiviridae* also belong to well supported monophyletic group suggesting that all four genes were also present in the common viral ancestor of the *Mimiviridae*. The *Mimiviridae* resolved into three clades that generally correspond to those previously described in the *Mimiviridae* (Claverie and Abergel 2018).

**Figure 4:**
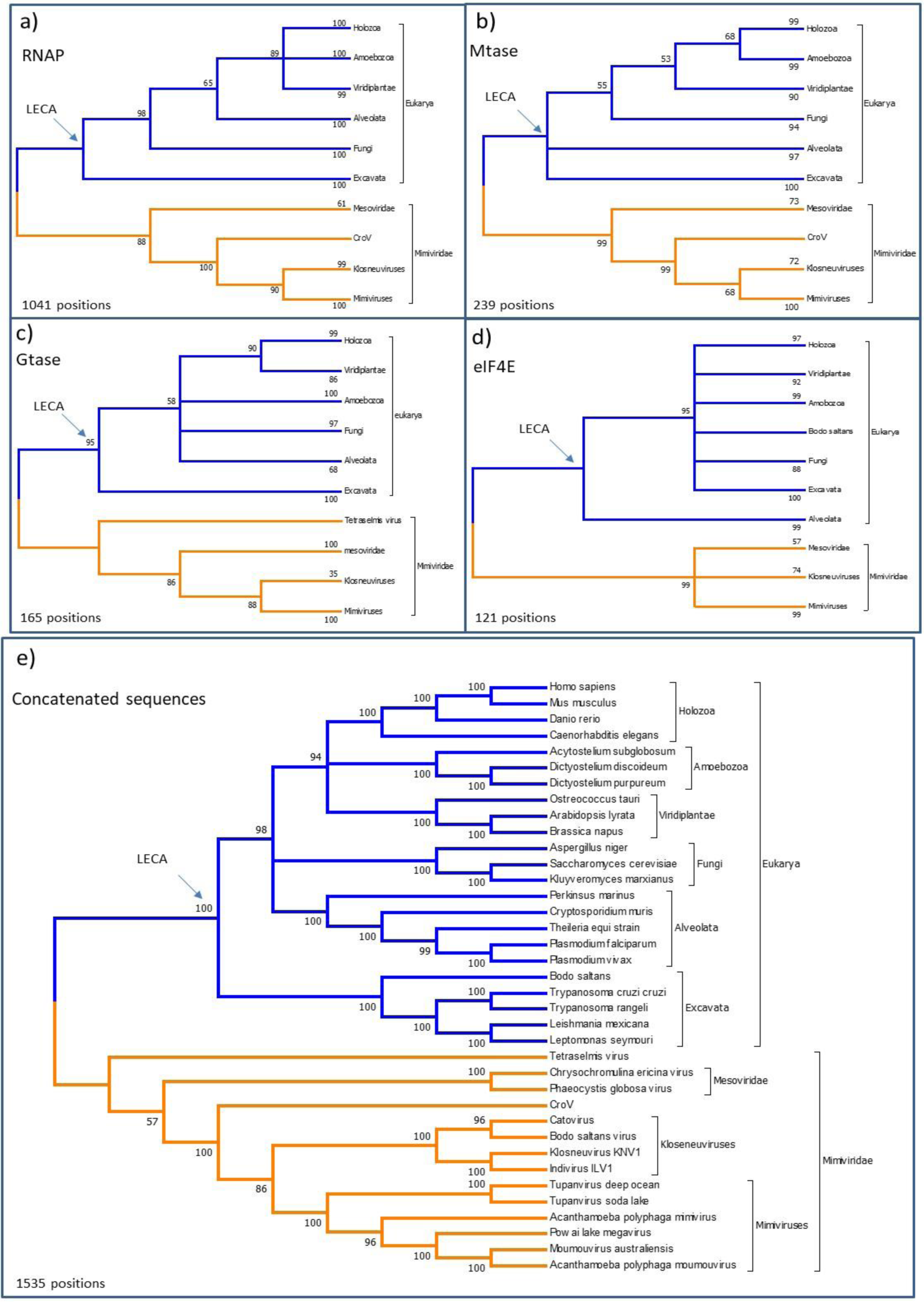
Unrooted phylogenetic trees of the mRNA capping pathway in selected eukaryotes and *Mimiviridae*. All five trees use sequences from the same set of carefully selected organisms (see Materials and Methods) and the proposed position of LECA is marked in each tree. The number of conserved amino acids in the final alignment for each gene is marked on the diagram. Trees were constructed and drawn using the ML method using default settings in MEGA7 with 1000 bootstrap replicates. NCBI accession numbers are given for each sequence in the Materials and Methods. *Mimiviridae* informal grouping names are based on Claverie and Abergel 2018. a) RNAP largest subunit gene tree. b) GTase gene tree. c) MTase gene tree. d) eIF4E gene tree e) Phylogenetic tree inferred from concatenation of all four gene sequences.

### The eukaryotic RNAP-II dedicated to capping mRNA shares a common ancestor with the *Mimiviridae* RNAP, and the common ancestor predates the origin of LECA

Although all the phylogenetic trees in **Figure 4** have been drawn with a root between the viral and eukaryotic versions of the genes, establishing the root of the MTase, GTase and eIF4E phylogenetic trees is challenging since the capping apparatus is unique to the eukaryotic domain. Thus only paralogues of these three genes exist outside the eukaryotes and the NCLDV viruses making it difficult to establish informative outgroups. In addition, despite being conserved, these genes are short and thus possess relatively little phylogenetic information. By contrast, the RNAP gene is a large phylogenetically informative gene that is found in all cellular domains. Since independent fossil evidence suggests that domain Archaea existed some two billion years before the appearance of LECA (Knoll 2015), and the eukaryotes apparently descend from a particular branch of the archaea (Spang et al. 2015), the RNAP large subunit is a suitable outgroup that can polarise the relationship between the eukaryotic RNAP-II and Mimiviral RNA polymerases. An additional advantage of the RNAP based tree is that all eukaryotes possess multiple RNAP’s (Sentenac 1985). Since these multiple RNAP’s were present in LECA, these can be used in concert with the archaeal sequences to firmly establish the root of the RNAP tree. Since both logic and the phylogenetic analysis performed here show that the RNAP, GTase, MTase and eIF4E genes are part of a co-evolving module responsible for producing and translating capped mRNA, it can be argued that establishing the root of the RNAP tree can be used to deduce the phylogeny of the entire capping apparatus.

As shown in **Figure 5**, using the archaeal RNAP subunit A’ and the homologous region of the eukaryotic RNAP-III to polarise the relationship between RNAP-II and the *Mimiviridae* RNAP shows that both the eukaryotic and Mimiviral genes descend from a common ancestral gene that predated the origin of LECA. The high bootstrap values give confidence that there is significant phylogenetic information in the alignment. In addition, both subtrees of the eukaryotic RNAP genes recapitulate the expected phylogenetic relationships between the eukaryotes, including the establishing the *Excavata* as the most divergent eukaryotic supergroup (Hampl et al. 2009). Furthermore, within the eukaryotic domains, all the chosen eukaryotes were assigned to their accepted branches. A parsimonious explanation of the observed tree is that the ability to produce m7G capped mRNA was a feature of the ancestor of both the eukaryotic RNAP-II and *Mimiviridae* RNA polymerase since both the eukaryotic and viral genes produce capped mRNA, whilst neither RNAP-III nor the Archaeal RNAP is associated with producing capped mRNA. Although other interpretations may be possible, the tree is entirely consistent with descent of the eukaryotic nucleus and the *Mimiviridae* from an ancient viral factory that could produce capped mRNA, a defining, core component of the apparatus required to uncouple transcription from translation by the eukaryotic nucleus that has not been observed in the archaeal relatives of the eukaryotes.

**Figure 5:**
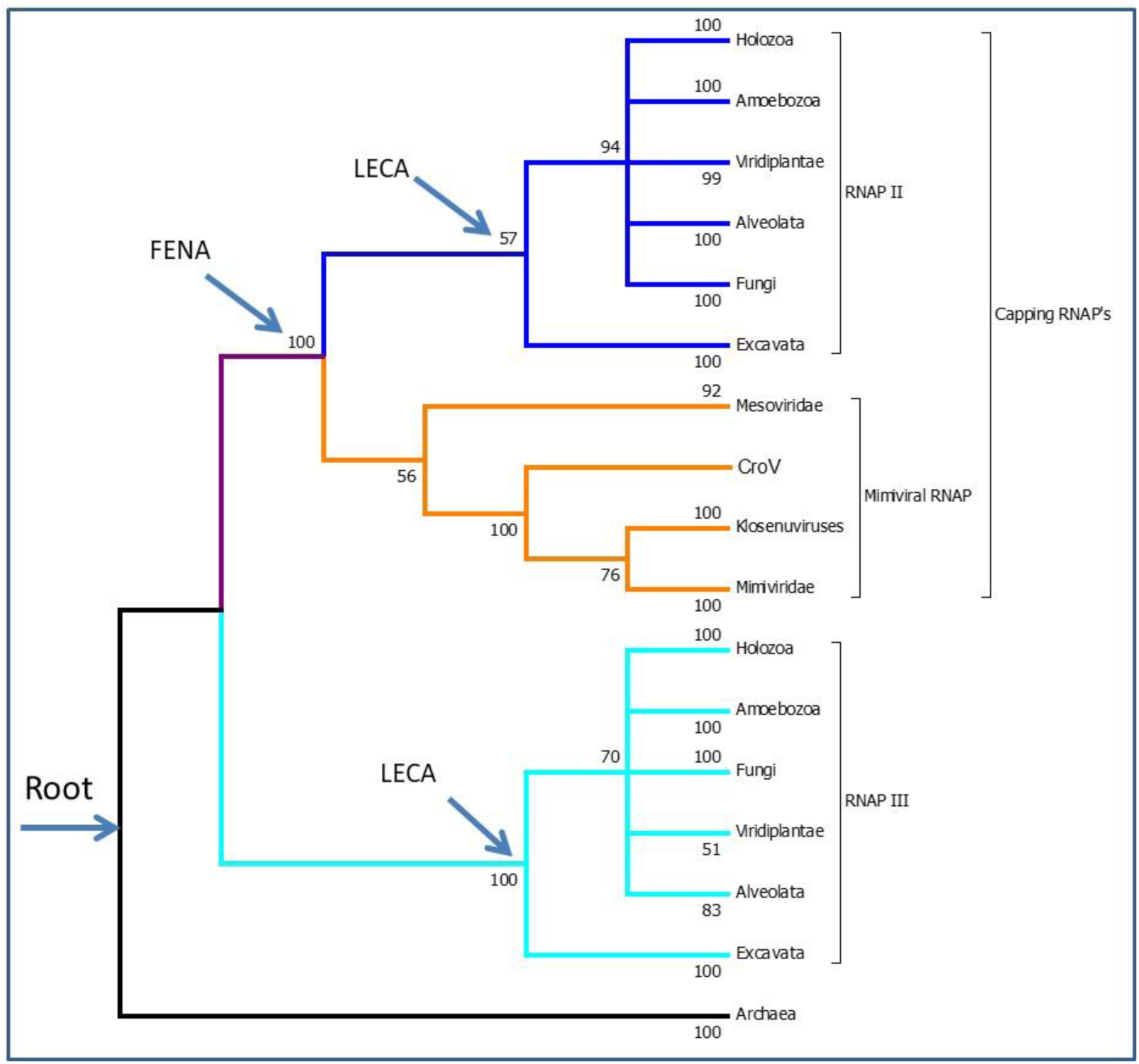
Maximum Likelihood tree of RNA polymerases using Archaeal RNAP subunit A’ as an outgroup. The RNAP A’ subunit of archaea was used as an out-group to establish the root of the largest subunit of the Mimiviral RNAP and Eukaryotic RNAP-II and RNAP-III genes. RNAP-II and RNAP-III are found to belong to two separate monophyletic groups. Both the RNAP-II and RNAP-III trees are robust, appropriately assign eukaryotes to their correct phylogenetic branches and re-capitulate the expected phylogenetic relationships between the eukaryotes including the early divergence of the *Excavata* (Hampl et al. 2009). The *Mimiviridae* tree is consistent with previous phylogenetic analyses of the *Mimiviridae* (Claverie and Abergel, 2018). This tree shows that the *Mimiviridae* and eukaryotic RNAP-II genes share a common ancestor. This ancestor existed before LECA and is consistent with the proposal that both descend from FENA, a proposed viral ancestor of both the Mimiviridae and the eukaryotic nucleus that infected an archaeal ancestor of the eukaryotes. Since both viral and eukaryotic RNAP-II synthesise m7G capped mRNA it can be inferred that the common RNA polymerase ancestor also produced capped mRNA. This tree was produced from an alignment of 64 sequences and 598 positions using Maximum Likelihood method and the JTT substitution model. Bootstrap values are indicated on each branch and are based on 1000 replicates. The tree and the computations were performed using MEGA7. NCBI accession numbers are given for each sequence in the Materials and Methods

## Discussion

Here it is shown that the apparatus used by eukaryotic nuclei to produce and translate capped mRNA is not found in the closest archaeal relatives of eukaryotes. This is significant since in the eukaryotic nucleus, the uncoupling of transcription from translation requires a complex highly evolved pathway consisting of hundreds of genes acting in concert (**Figure 2**) and the m7G cap is critical to this pathway since it is used to prime mRNA for processing, nuclear export, and cytoplasmic translation (**Figure 2**). The absence of the m7G apparatus implies that the highly complex pathway for uncoupling transcription from translation is also absent from archaeal relatives of the eukaryotes. This presents a major biological paradox since such a complex pathway incorporating the concerted action of hundreds of genes unique to the eukaryotic domain implies a long evolutionary history, yet no sign of the pathway is found in the closest archaeal relatives of the eukaryotes.

Although the appartus for producing capped mRNA is absent from the archaeal relatives of the eukaryotes, the apparatus is present in the *Mimiviridae* which is consistent with the postulates of the VE hypothesis. Phylogenetic analysis performed here demonstrates that viral and eukaryotic genes form discrete monophyletic clades, and that both viral and eukaryotic clades descend from a common ancestor that existed prior to the appearance of LECA. This pattern is consistent with proposal that the eukaryotic nucleus and the *Mimiviridae* both descend from a First Eukaryotic Nuclear Ancestor (FENA).

Prior to the discovery of the nucleus-like viral factory of *201 Φ2-1*, the ability to uncouple transcription from translation was thought to be an exclusive innovation of the eukaryotic nucleus. Thus, arguments could be made that the viral factory of the Mimiviruses had evolved by borrowing genes from the nucleus to allow it to establish the eukaryotic uncoupling of transcription from translation. However, since *201 Φ2-1* infects bacteria it seems very unlikely that it obtained its ability to build a viral factory and uncouple transcription from translation from the eukaryotes, but rather indicates this ability has evolved in prokaryotic viruses as part of their replication cycle. Thus the discovery of *201 Φ2-1* demonstrates that uncoupling of transcription from translation is most likely a viral innovation, and since prokaryotes existed billions of years before the origin of the eukaryotes, viral factories potentially existed for billions of years before the origin of LECA. Studies on a relative of phage *201 Φ2-1* (*PhiKZ*), show the viral factory appears to shield phage DNA from host immune systems including the CrispR cas system (Mendoza et al. 2018). Thus viral factories may have evolved to provide biological protection from various anti-phage systems possessed by prokaryotic hosts (Hendrickson and Poole 2018).

The modern nucleus is clearly differentiated from any member of the *Mimiviridae* by its ability to construct a fully functional translational system including ribosomes. In the absence of their own translational machinery, all known viruses are dependent upon their host’s translation machinery to produce polypeptides required for their own reproduction. Thus all mRNAs produced by viruses accordingly engage cellular ribosomes to ensure translation (Jan et al. 2016). However, when the *Mimivirus* was first discovered the “most unexpected discovery was the presence of numerous genes encoding central protein-translation components” (Raoult et al. 2004). The discovery of the *Klosneuvirus* increased the number of translation related genes found in viruses to levels that far exceeds that seen in the original *Mimivirus* (Schulz et al. 2017) and sequencing of the *Tupanvirus* genome revealed that some members of the *Mimiviridae* possess a translation associated gene set that ‘only lacks the ribosome’ (Abrahão et al. 2018). Amongst this set of translational genes is up to 70 tRNA, 20 aaRS, 11 factors for all translation steps and factors related to tRNA/mRNA maturation and ribosome protein modification (Abrahão et al. 2018). Since it appears that the ancestor of the *Mimiviridae* did not possess all these functions, the appearance of so many translation related genes in viruses such as the *Kloseneuvirus* and the *Tupanvirus* suggests that they acquired these components of the eukaryotic translational machinery via a piecemeal capture process (Schultz et al. 2017).

The expanded viral repertoire of translational genes found in the *Tupanvirus* and *Klosneuvirinae* suggest that there is selective pressure to acquire these genes in some branches of the *Mimiviridae*. This capture process may also be acting amongst the modern giant phage where captured ribosomal genes appear to be part of the mechanism(s) by which phage direct the host translation apparatus to selectively translate viral mRNA (Al-Shayeh et al. 2019). If the VE hypothesis is valid and a similar capture process was operating before the origin of the first eukaryotes, the translational apparatus acquired by FENA could only have been captured from prokaryotic cells. Since FENA is proposed to have infected an Asgardian ancestor, many of the translation related genes would be derived from its archaeal Asgardian host and directed to enhancing translation of the viral transcripts by the host’s archaeal translational system. Consistent with this proposal, it is known that eukaryotic nuclei possess a core set of archaeal related translation initiation factors including eIF1A, eIF2, eIF2B, eIF4A, eIF5B and eIF6 (Jagus et al. 2012), and a core set of eukaryotic specific initiation factors (eIF5, eIF4E, eIF4G, eIF4B, eIF4H and eIF3) (Jagus et al. 2012). With the exception of eIF5, all these eukaryotic specific initiation factors are involved with 5’-cap-binding and scanning processes required for translation of capped eukaryotic mRNA (Jagus et al. 2012). Furthermore in the process of evolving into the nucleus, a viral ancestor of the nucleus must have acquired the ability to synthesise uncapped rRNA and tRNA, and thus a part of the transition into a fully autonomous nucleus would have been the capture by the virus of second and third RNA polymerase dedicated to the synthesis of non-capped RNA associated with functioning of the ribosomes.

If the VE hypothesis can be accepted, the descent of the nucleus from a viral factory provides a plausible resolution to several of the major paradoxes associated with the origin of the nucleus. That is, if the nucleus descends from a viral factory and the viral factory set up by FENA was similar in structure and function to the *201 Φ2-1* and *Mimiviridae* viral factories (**Figure 3**), the VE hypothesis explains why the nucleus is mainly an information containing and processing compartment, why it’s boundary selectively controls the entry and exit of proteins and nucleic acids, why it exports mRNA into the cytoplasm, why it contains no functional ribosomes, why it possesses linear rather than circular chromosomes, why it is positioned in the cell by the tubulin cytoskeleton, and as explored in this paper, why the eukaryotes possess highly evolved complex machinery to allow uncoupling of transcription from translation with no prokaryotic precedents. It also provides a rationale for the neo-functionalisation of RNA polymerases in the eukaryotes since the viral factory introduces its own RNA polymerase specifically dedicated to the transcription of capped viral mRNA destined for translation in the cytoplasm. The origin of the nucleus from a viral ancestry has also been shown to provide a plausible mechanistic model for the origin of mitosis, meiosis and the sexual cycle (Bell 2006, Bell 2013), a problem described as the queen of evolutionary problems (Bell 1982). Thus the origin of the nucleus from a viral factory addresses many of the challenges required to explain the apparently abrupt appearance of a fully formed and functional nucleus in LECA, despite its complete absence from bona-fide archaeal relatives such as members of the Asgard archaea.

It should be noted that the VE hypothesis is not a pure ‘endosymbiotic theory’. According to the VE hypothesis (Bell 2001), the eukaryotic cell is descended from an archaeal ancestor of the eukaryotic cytoplasm, a bacterial ancestor of the mitochondrion, and as explored in this paper, a viral ancestor of the nucleus. Although the archaeal ancestor of the cytoplasm may had a mutually beneficial symbiotic relationship with a bacterium leading to the origin of the mitochondria, the host archaeon did not gain any benefit from the viral infection, rather the archaeon host was enslaved by the virus and its genome was ultimately destroyed.

However, like the endosymbiotic theories for the origin of the mitochondria and the chloroplasts, the VE hypothesis deals with complex irreversible events that are difficult to directly test (Margulis 1975). In the case of the mitochondria, it took nearly 100 years before the consilience of evidence built up sufficiently for the endosymbiotic origin the mitochondria to become (almost) universally accepted. Although a more radical concept than endosymbiosis, if the VE hypothesis is similarly supported by the accumulation of multiple lines of evidence, it will introduce a major paradigm shift in our understanding of the evolution of complex life on earth. In particular, if the VE hypothesis is ultimately accepted, it implies that the eukaryotic cell derives from a consortium of three organisms that became integrated to such an extent that they created an emergent ‘super-organism’. The novel features of this emergent ‘super-organism’ allowed it to escape the limitations of prokaryotic evolution and evolve to levels of unprecedented organismal complexity.

## Materials and Methods

### Choice of eukaryotic organisms

#### i) Eukaryotes

The organisms used in this study were carefully selected to cover all the relevant groupings of eukaryotes, whilst limiting the complexity of the phylogenetic analysis. Currently 5 or 6 eukaryotic supergroups are proposed to cover the vast majority of eukaryotic diversity (Hampl et al. 2009). The present study focussed on ‘model’ organisms for the phylogenetic trees so that there was significant knowledge of their molecular biology of at least one or more of the divisions. To represent the *Holozoa, Homo sapiens, Mus musculus, Danio rerio* and *Caenorhabditis elegans* were chosen since each is a model organism, and the phylogenetic relationships are well established. To represent *Amoebozoa, Dictyostelium disocoidium* was chosen since it is a model organism. *Dictyostellium purpurem* and *Acytostelium subglosum* were chosen as suitably distant relatives. To represent the Fungi, *Saccharomyces cerevisiae, Kluyveromyces marxianus* and *Aspergillus niger* were chosen since all three are model organism and the phylogenetic relationships are well understood. To represent *Viridiplantae, Arabidopsis lyrata* was chosen as a model species and *Brassica napus* was chosen as a relatively close relative. *Ostreococcus tauri* was chosen as a distant algal relative of the land plants. To represent the SAR group, focus was placed on the *Alveolata* group since members such as *Plasmodium* and *Cryptosporidium* have been studied in depth at a molecular level. To ensure the robustness of the tree and limit the effects of long-branch attraction, *Plasmodium falciparum* and *Plasmodium vivax* were chosen as close relatives whilst *Theileria equi* strain WA, *Cryptosporidium muris*, and *Perkinsus marinus* were chosen as increasingly distantly related members of the *Alveolata*. To represent the *Excavata*, members of the *Trypanosoma* were chosen since they are model organisms that have been studied in depth at molecular level. To ensure robustness of the tree and to minimise the effects of long-branch attraction, *Trypanosoma cruzi cruzi* and *Trypanosoma rangeli* were chosen as close relatives whilst *Leishmania mexicana, Leptomonas seymouri* and *Bodo saltans* were chosen as increasingly distantly related members of the *Excavata*. The organisms listed above include members of all 5 or 6 major clades. In addition, complete genomes are available for each of the organisms listed, ensuring that the phylogenetic trees included exactly the same organisms.

#### ii) Mimiviridae

Only members of the *Mimiviridae* containing clear homologues to RNAP, GTase, MTase and eIF4E were chosen for analysis. Based on phylogenetic analysis by Claverie and Abergel, 2018, the following viruses were chosen to represent three informal groupings of the *Mimiviridae*. *Mesomimivirinae*: *Tetraselmis* virus, *Chrysochromomulina ericina* virus and *Phaeocystis globosa* virus. *Klosnuevirinae: Klosneuvirus, Catovirus, Indivirus* and *Bodo saltans* virus. *Megavirinae*: *Acanthamoeba polyphaga mimivirus, Powai lake megavirus, Moumouvirus australiensis, Acanthamoeba polyphaga moumouvirus, Tupanvirus deep ocean* and *Tupanvirus soda lake. Cafeteria roenbergensis* virus (CroV) is basal to the *Klosenuvirinae* and *Megavirinae* and does not appear to have other close relatives available yet.

### Choice of sequences

#### i) RNAP subunits

Of the three RNA polymerases in eukaryotes, the RNAP-II is the one intimately associated with the capping of transcripts. The largest RNAP-II subunit possesses a carboxy terminal domain (CTD) consisting of a heptapeptide repeat region that is involved in mRNA processing including capping, splicing and polyadenylation (McCracken et al. 1997). Homologues of RPO21, the largest subunit of RNAP-II of *S. cerevisiae* were identified. With the exception of the members of the *Excavata* and *Perkinus marinus*, a CTD heptapeptide repeat region was readily identified in all RNAP-II subunits used in the phylogenetic analysis. Although the heptapeptide repeat is absent from the *Excavata* studied, *Trypanosoma* RNAP-II genes possesses a non-canonical C-terminal extension (Smith et al. 1989). As a result, the *Trypanosoma cruzi cruzi* RNAP-II was used to identify RNAP-II homologues in the *Excavata* clade. In the *Mimiviridae* only one homologue of the largest subunit of RNAP-II was detected.

#### ii) GTase and MTase

Although three enzymatic functions are universally required to produced capped mRNA (Kyrieleis et al. 2014), only the GTase and MTase are monophyletic in eukaryotes, with the TPase apparently originating from two independent sources (Ramanathan et al. 2016; Kyrieleis et al. 2014). In *S. cerevisiae* and most other unicellular eukaryotes such as *Alveolata* all three functions are encoded by separate genes. In both *Holozoa* and *Viridiplantae* the TPase and GTase are encoded in the same polypeptide. In *Excavata*, two capping complexes are present (Takagi et al. 2007). Of these, the gene encoding both the GTase and MTase in the same polypeptide is essential for growth and adding the m7G cap and was thus chosen for phylogenetic analysis. In the *Mimiviridae*, all three functions are present in the same polypeptide.

#### iii) eIF4E

Although in the yeast *Saccharomyces cerevisiae*, there is only one eIF4E gene, the core role of eIF4E in protein translation has meant that in higher eukaryotes several paralogous eIF4E genes have evolved that encode distinctly featured proteins. In addition to regular translation initiation, these paralogues are involved in the preferential translation of particular mRNAs or are tissue and/or developmental stage specific. For example, eight such genes have been found in *Drosophila* and five in *Caenorhabditis* (Frydryskova et al. 2018). In humans where there are multiple paralogues, the three isoforms of eIF4E1 bring the mRNAs to the ribosome via an interaction with scaffold protein eIF4G (Frydryskova et al. 2018). As a result, in this study the human eIF4E1 isoform was used to conduct blast searches of *Holozoa*, and the hits with the highest blast score were taken for phylogenetic analysis. In *Arabidopsis*, the EIF4E1 is expressed in all tissues except in the cells of the specialization zone of the roots whereas the At. EIF4E2 mRNA is particularly abundant in floral organs and in young developing tissues (Rodriguez et al. 1998). The Arabidopsis EIF4E1 gene was thus used in blast searches of plants, and the genes with the highest homology taken for phylogenetic analysis. Where molecular knowledge was insufficient for such rational sequence selection, the homologue with the highest homology to the *Saccharomyces* gene was identified, and provided that the gene possessed regions equivalent to the structurally important regions that bind to the m7G cap (Marcotrigiano et al. 1997), this gene was used to identify the closest homologues within the supergroup. With the exception of the Tupanviruses, the *Mimiviridae* were found to encode only one eIF4E homologue. In the case of the Tupanvirus, two eIF4E homologues were identified. In this case, only one of the homologues was included in the phylogenetic analysis. The homologue with the highest homology to the Mimivirus homologue was used in both cases.

### Phylogenetic analysis

Homology searches were carried out using the BLASTp and psi-BLAST algorithms (Altschul et al. 1997). MEGA7 (Kumar et al. 2016) was used for all phylogenetic analysis. Unless otherwise stated all program parameters for homology searching and domain identification were left at their respective defaults. Protein alignments were performed using MUSCLE. Once alignments were completed for all organisms for a particular alignment, the alignments were trimmed and used for tree construction. The evolutionary histories were inferred by using the Maximum Likelihood method based on the JTT matrix-based model (Jones et al. 1992). All bootstrap consensus trees were inferred from 1000 replicates (Felsenstein 1987) and is taken to represent the evolutionary history of the taxa analyzed (Felsenstein 1987). Branches corresponding to partitions reproduced in less than 51% bootstrap replicates were collapsed. The percentage of replicate trees in which the associated taxa clustered together in the bootstrap test (1000 replicates) are shown next to the branches (Felsenstein 1987). Initial tree(s) for the heuristic search were obtained automatically by applying Neighbor-Join and BioNJ algorithms to a matrix of pairwise distances estimated using a JTT model, and then selecting the topology with superior log likelihood value.

